# SNP Variable Selection by Generalized Graph Domination

**DOI:** 10.1101/396085

**Authors:** Shuzhen Sun, Zhuqi Miao, Blaise Ratcliffe, Polly Campbell, Bret Pasch, Yousry A. El-Kassaby, Balabhaskar Balasundaram, Charles Chen

**Affiliations:** Department of Biochemistry and Molecular Biology, Oklahoma State University, Stillwater, OK 74078; Department of Forest and Conservation Sciences, Faculty of Forestry, The University of British Columbia, Vancouver, B.C. Canada, V6T 1Z4; Center for Health Systems Innovation, Oklahoma State University, Stillwater, OK 74078; Department of Integrative Biology, Oklahoma State University, Stillwater, OK 74078; Department of Biological Sciences, Northern Arizona University, Flagstaff, AZ 86011; School of Industrial Engineering and Management, Oklahoma State University, Stillwater, OK 74078

## Abstract

High-throughput sequencing technology has revolutionized both medical and biological research by generating exceedingly large numbers of genetic variants. The resulting datasets share a number of common characteristics that might lead to poor generalization capacity. Concerns include noise accumulated due to the large number of predictors, sparse information regarding the *p* ≫ *n* problem, and overfitting and model mis-identification resulting from spurious collinearity. Additionally, complex correlation patterns are present among variables. As a consequence, reliable variable selection techniques play a pivotal role in predictive analysis, generalization capability, and robustness in clustering, as well as interpretability of the derived models.

*K*-dominating set, a parameterized graph-theoretic generalization model, was used to model SNP (single nucleotide polymorphism) data as a similarity network and searched for representative SNP variables. In particular, each SNP was represented as a vertex in the graph, (dis)similarity measures such as correlation coefficients or pairwise linkage disequilibrium were estimated to describe the relationship between each pair of SNPs; a pair of vertices are adjacent, i.e. joined by an edge, if the pairwise similarity measure exceeds a user-specified threshold. A minimum *K*-dominating set in the SNP graph was then made as the smallest subset such that every SNP that is excluded from the subset has at least *k* neighbors in the selected ones. The strength of

*k*-dominating set selection in identifying independent variables, and in culling representative variables that are highly correlated with others, was demonstrated by a simulated dataset. The advantages of *k*-dominating set variable selection were also illustrated in two applications: pedigree reconstruction using SNP profiles of 1,372 Douglas-fir trees, and species delineation for 226 grasshopper mouse samples. A C++ source code that implements SNP-SELECT and uses Gurobi™ optimization solver for the *k*-dominating set variable selection is available (https://github.com/transgenomicsosu/SNP-SELECT).

## Introduction

With the rapid advancement of DNA sequencing technology, the volume and dimension of biological and medical data have been increasing at an unprecedented rate. Accompanying such high volume genetic data, the ‘curse of dimensionality’ has not only hindered genetic discovery by the demanding computing resource required; model noise also accumulates due to a large number of predictors. The spurious collinearity of the large number of predictors that causes over-fitting and mis-identification of models raises serious concerns. As a result, variable selection methods are playing pivotal role in predictive analyses, clustering, and classification. For example, Song et al. (1) showed that with a selected subset of 1,500 SNP, comparable predictability for traits like grain yield could be achieved without using the entire SNP dataset (>10K SNPs). In a closely related wheat population, predictability obtained using 34,749 SNPs could be attainable with as few as 1,827 SNPs (2). Research in a bi-parental bread wheat population genotyped with 485 SNP markers showed prediction accuracy plateaued with 128-256 markers, beyond which accuracy started to decline (3). A similar benefit from SNP variable selection for prediction is also seen in dairy cattle and maize breeding programs (4, 5). In a genomic prediction study, where predictability of human height, high-density lipoprotein cholesterol (HDL) and body mass index (BMI) was examined for approximately 3,000 individuals, supervised variable selection was recommended in the GBLUP (genomic best linear unbiased prediction) framework (Bermingham et al. (6)).

Crucial to both analyzing and interpreting high dimensional datasets, significant effort has been directed towards exploring variable selection processes by removing features that might be either redundant or irrelevant to the problem, for better predictability, or computational efficiency and informativeness (7). This effort includes the logistic regression method (8), the penalized regression method (9-11), partial least squares regression (PLSR) (12), sure independence screening strategy (13), multi-stage regression methods (14), sorted l-one penalized estimation (SLOPE) via convex optimization (15), recurrent relative variable importance measure (r2VIM) (16), to name a few. However, these methods were designed to reduce variables from a statistical perspective in order to ease the process of prediction or assist GWAS (genome-wide association study) analysis, in which knowledge of phenotypic data is required.

In the era of population genomics (17), many *Fst*-based genome-scan methods utilize large datasets such as SNP chips or genome complexity reduction approaches like RAD tags (18) and genotyping-by-sequencing (GBS) (19, 20), to estimate genetic parameters (21). Identifying adaptive evolution and candidate genomic regions under selection is increasingly feasible, thanks to the development of sophisticated analytical tools for genome-scale polymorphism data (22-25). Given the data volume, most of these Bayesian approaches suffer from extended computational time requirement (21) due to tedious numerical approximation procedures like Markov chain Monte Carlo (MCMC) (24) or reverse jump (RJ)-MCMC (23). Furthermore, accurate inference of demographic parameters such as effective population sizes, migration rates, and divergence times between populations depends largely on the use of neutral marker data (26-28). In other words, SNP variable selection methods without the use of phenotypic data are desirable for the purpose of reducing the bias caused by confounding variables, for minimizing computational load, and for avoiding the potential problem of allele frequency correlations in, for example, the Lewontin and Krakauer (LK) test (21, 29).

In this paper, we present SNP-SELECT, a variable selection algorithm based on a graph-theoretic approach that uses generalized dominating sets for a large volume of SNP data without the use of phenotypes. Application of graph theory to variable selection or data reduction has been seen in many data mining applications (30-32). Typically, this involves clustering the data points into groups and using one point to represent each cluster, from which the network clustering (33) procedure would derive a much smaller number of clusters, resulting in variable selection. In our cases, data points (SNPs) are represented by vertices and an edge exists if two data points (two SNPs) are similar or related in a certain way (i.e., in linkage disequilibrium (LD) or in correlation). We show the use of LD with an example; it is, however, important to note that the similarity criterion used to construct the network model can be based on any relationship measurement. The advantage and robustness of SNP-SELECT is also demonstrated with simulated datasets, and with empirical datasets for a Douglas-fir (*Pseudotsuga menziesii*) breeding population and for populations of three grasshopper mouse species (*Onychomys* spp.).

## Material and Methods

### Generalized Graph Domination

Let *G* = (*V*, *E*) be a graph with vertex set *V* and edge set *E* ⊆ [*V*]^2^ (see (34) for basic graph theory concepts and notations). The *open neighborhood* of a vertex *v* is the set *N*(*v*) of vertices adjacent to vertex *v*. Note that *v* ∉ *N* (*v*) and the *closed neighborhood* of vertex *v* is denoted by *N*[*v*] = {*v*} ⋃ *N* (*v*).

**Definition 1 (35)** Given a positive integer *k* and a graph *G* = (*V*, *E*), a subset of vertices *D* is said to be *k-*dominating if | *N* (*v*) ⋂ *D* | ≥ *k* for every vertex *v* ∉ *D*.

If *D* is a *k-dominating* set, then every vertex in *V − D* is said to be *k-*dominated. A *minimum k-dominating set* is one of smallest cardinality in the graph and this cardinality is called the *k-domination number* of the graph, denoted as *γ*_*k*_(*G*). Note that the *k*-domination number of a graph increases as parameter *k* increases and the model becomes more restrictive as more neighbors are needed for each vertex outside the set to be *k*-dominated. Hence, every 2-dominating set is also a 1-dominating set, but the converse is not true. Intuitively, as the parameter *k* increases, we expect the *k*-dominating set to be a more reliable representation of the dataset as each point has at least *k* similar points in the *k*-dominating set. Hence, the choice of *k* must balance two conflicting criteria: solution fidelity (how well the dataset is represented) and solution size (how many data points are selected). To illustrate, graphic presentations of *k*-dominating sets for *k* = 1 and *k* = 2 were showed in Figure 1(a); and Figure 1(b) illuminated 1-dominating set using neural network data for the nematode, *C. elegans* (36, 37). Neurons are represented by vertices in this neural network and as long as two neurons communicate with each other, an edge exists between them. The big dots in Figure 1(b) mark a 1-dominating set, and all the small dots (vertices) have at least one neighbor of the same color, which identifies the cluster.

**Fig 1.**
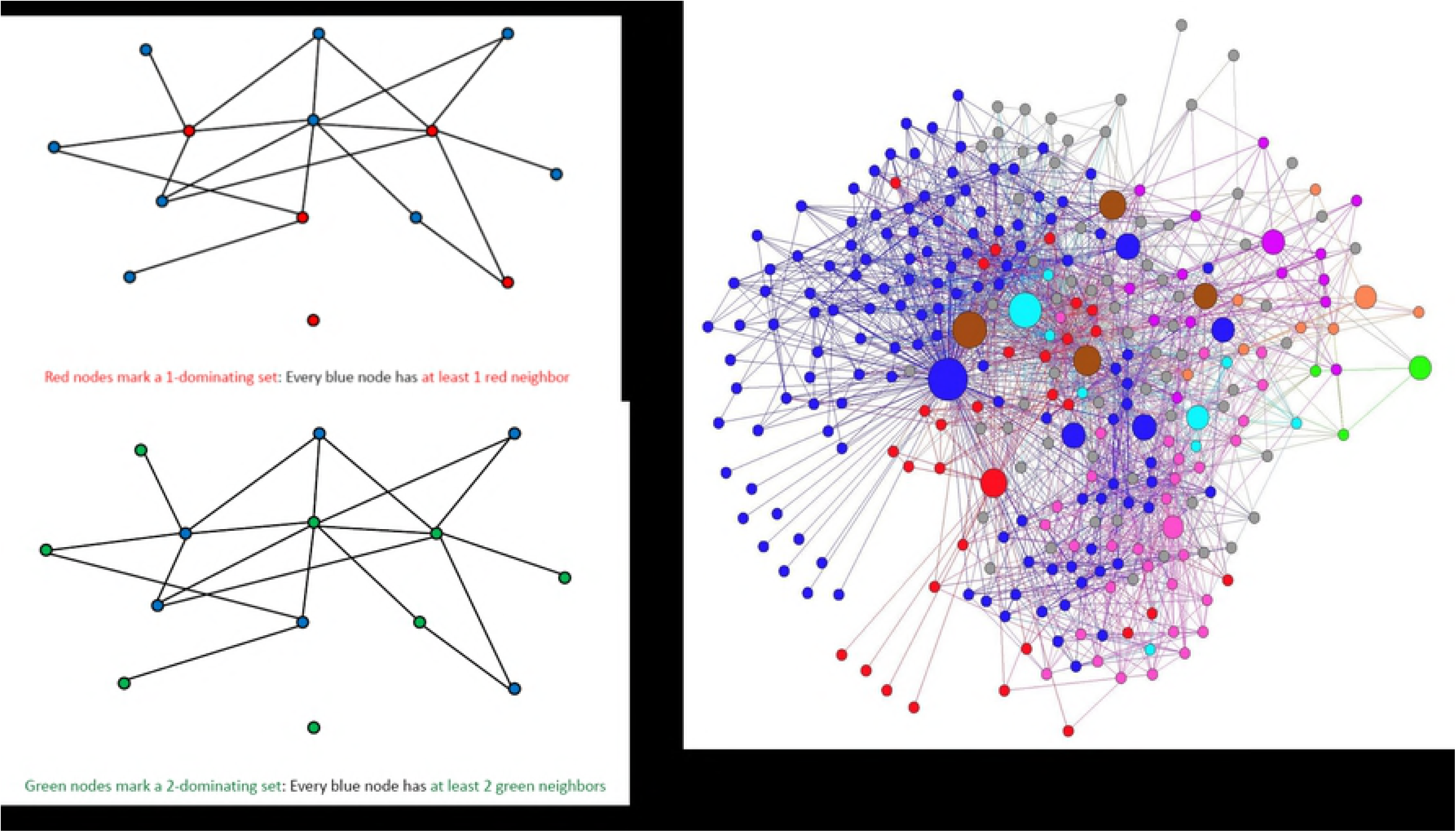
(a) Illustration of 1-dominating set and 2-dominating set; (b) Illustration of 1-dominating set using the neural network data of *C. elegans* (36, 37): the big nodes mark a 1-dominating set, and all the small nodes have at least 1 same color neighbor.

Clustering a graph via *k*-dominating sets, especially with *k* = 1, is a popular technique in telecommunication and wireless networks (38). If *D* is a 1*-dominating* set, then for each vertex *v* ∈ *D* the closed neighborhood *N*[*v*] forms a cluster that altogether cover *V*. Since by definition, every vertex not in the 1-dominating set has a neighbor in it and hence, is assigned to at least one cluster. Since the problem of finding a minimum *k*-dominating set is NP-hard (39), heuristic approaches and approximation algorithms have been proposed to find a small *k*-dominating set in the given graph (40). However, the approach employed in this article to solve this combinatorial optimization problem was to formulate it as an integer program (41), implement and solve it using a state-of-the-art solver that employs a branch-and-cut algorithm with built-in primal heuristics and other presolve reductions among others. Given a positive integer *k* and a graph *G* = (*V*, *E*), the problem of finding a minimum *k*-dominating set can be formulated as the following linear integer program in binary variables.

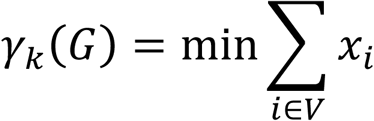

subject to:

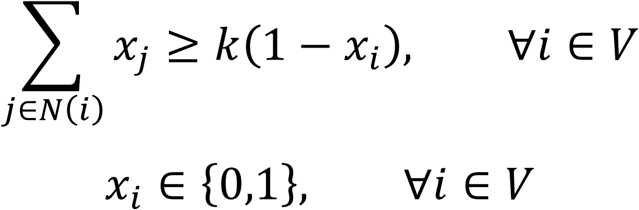

In any feasible solution *x* to this formulation, the binary variable *x*_*i*_ = 1 if and only if vertex *i* is included in the *k*-dominating set *D*, which is given by *D* = {*i* ∈ *V*: *x*_*i*_ = 1}. The constraints ensure that if a vertex *i* is excluded from the *k*-dominating set *D,* i.e. *x*_*i*_ = 0, at least *k* of its neighbors must be included.

### Pairwise Relationship between SNPs

The pairwise relationship (similarity or distance) between SNP variables primarily determines the structure of the graph *G*, and different ways for quantifying the pairwise relationship can influence the structure of the graph *G*, especially the sets of edges. Currently, many methods exist to measure the pairwise relationship of SNPs, for example, Hamming distance (42), mutual information (43, 44), allele sharing index (45, 46), and linkage disequilibrium (LD) (47-49), to name a few. We chose to use the frequently used LD approach to describe the pairwise relationship between SNP variables in this study, although the proposed approach continues to work with other similarity measures as well. The square of correlation coefficients (*r*^2^) for SNP variables were calculated to represent the values in the LD matrix (refer to (50) for the details). Since haplotype frequencies for each pair of SNPs are unknown, the expectation maximization algorithm (51) was applied to infer the haplotype frequencies in the LD calculation.

With a user-defined threshold (*λ*), an edge exists only if the pairwise relationship between the two SNPs (vertices) is greater than *λ*. Thus, for any given pairwise relationship measurement, as *λ* increases, the number of edges in the graph decreases, and consequently the number of isolated (independent) vertices in a graph can increases. For any positive integer *k*, an isolated vertex in the graph cannot be *k*-dominated by any other vertex, and must be included in any *k*-dominating set. In fact, this observation holds more generally for any vertex with fewer than *k* neighbors in the graph.

### Scheme of SNP-SELECT

The details of SNP-SELECT are summarized as follows:

**Step 1:** Construct a graph model *G* = (*V*, *E*): Let *V* be the set of all SNPs and *E* is initially empty;

**Step 2:** Calculate linkage disequilibrium *w*_*ij*_ for each pair of SNPs *i*, *j* ∈ *V*;

**Step 3:** An edge between SNPs *i* and *j* is created if *w*_*ij*_ > *λ*;

**Step 4**: Identify isolated SNPs *I* ← {*i* ∈ *V*: *N*(i) = Ø};

**Step 5:** Find a minimum *k*-dominating set in *G* − *I*.

All experiments/analyses reported in this article were conducted on a 64-bit Linux compute node of a high performance computing cluster with 96GB RAM and Intel Xeon E5620 2.40GHz processor. The algorithm was implemented using C++, and the integer programming formulation for the minimum *k* -dominating set problem was solved using the Gurobi^™^ optimizer 6.0 with default settings (52). Given a running time limit, Gurobi^™^ either returned an optimal solution, or a feasible solution with a gap to a lower bound on the optimal solution. Experiments/analyses reported in this study were performed with a 1-hour running time limit for Gurobi^™^. The solution returned by Gurobi^™^ was used to identify the representative subset of the original dataset.

In our preliminary analyses, we found that when *λ* is small, the graph model tends to be very dense with an extremely large number of edges. When several thousands of SNPs are involved, such graphs can exceed memory limits during computation and result in a memory crash, before a feasible solution can be derived. Also, very small thresholds may not necessarily be realistic to capture similarities between SNPs. To address this issue, a stepwise search was implemented in SNP-SELECT for large SNP datasets as follows:

**Step 1:** Construct a threshold set *T* = {*λ*_1_,*λ*_2_, *λ*_*L*_}, where *λ*_1_ > *λ*_2_ > > *λ*_*L*_, and *λ*_*L*_ is the desired threshold, *λ*_*L*_ ← *λ*, and *λ*_*h*_ − *λ*_*h*+1_ equals a predefined step; Let *h* = 1, and *V*_1_ be the set of all SNPs;

**Step 2:** Construct *G*_*h*_ on *V*_*h*_ using *λ*_*h*_;

**Step 3:** Identify isolated SNPs *I*_*h*_ ← {*i* ∈ *V*_*h*_: *N*(i) = Ø};

**Step 4:** Find a minimum (or a small) *k*-dominating set *S*_*h*_ in *G*_*h*_ − *I*_*h*_, let *Y*_*h*_ ← *S*_*h*_ ⋃ *I*_*h*_;

**Step 5:** If *h* = *L*, return *Y*_*h*_, STOP; else *V*_*h*+1_ ← *Y*_*h*_, *h* ← *h* + 1, go to **Step 2**.

In brief, this step-wise search of SNP-SELECT first finds a minimum *k*-dominating set *Y*_1_ (or the best solution available) on a graph model based on a larger threshold. Then the threshold is lowered to focus on the graph induced by *Y*_1_. The data size of current step is the output of previous step. This process is repeated until a desired low threshold is reached. The feature selection problem of large datasets is thus solved by iteratively reducing the value of threshold.

### Simulation Studies

To demonstrate the capacity of the *k*-dominating set algorithm to identify independent variables, and to select proxy variables among highly correlated ones, a simulated dataset that included 10 synthetic undirected networks with *n* = 1000 vertices were used to represent SNPs. In this synthetic network dataset, the pairwise relationships between SNPs (vertices), the weighted edge (*w*_*ij*_) between each pair of vertices (*i*, *j*), were generated using a uniform distribution over [0,1]. The randomly chosen edge weights, denoted by *a*_*l*_, where 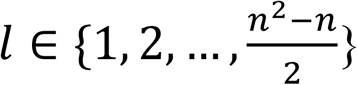, and without loss of generality assumed to be in increasing order, were assigned to edges using the following algorithm such that *w*_*i*, *j*_ < *w*_*i*, *j*+1_ and *w*_*i*, *j*_ < *w*_*i*+1,*j*_.

**Step 1:** Initialize *l* ← 1;

**Step 2: for *i* = 1** to *n* – 1

**Step 3: for *j* = *i* + 1** to *n*

**Step 3:** *w*_*ij*_ ← *a*_*l*_;

**Step 4:*l*** ← *l* + 1;

**Step 5**: **end-for**

**Step 6: end-for**

A correlative relationship among SNP variables, or linkage disequilibrium (LD), is the non-random association between SNP alleles. The distribution of these relationships among SNPs in a given genome tends to be greater when SNPs are closely located; this correlation diminishes quickly as genomic distance between SNPs gets larger, e.g. LD decay (53). As a result, the distribution of correlative relationships among SNPs is a mixture of a small number of highly correlated SNPs with a large number of SNPs in low correlations. Assigning edge weights in increasing order is a simple way to guarantee that only part of the vertices has low edge weights close to 0, which can be used to define the independent variables. Meanwhile, we can also identify a subset of vertices with edge weights higher than a predefined threshold within this set, which could be used to define the independent variables and highly correlated variables.

A vertex *i* that has all neighbors with *w*_*ij*_ < 0.1, where *j* ∈ *V* and *j* ≠ *i*, was defined as an independent variable. The subset generated by SNP-SELECT has to include all the independent variables to confirm that the *k*-dominating set based approach is able to identify independent variables. Highly correlated variables were defined as a subset (*P*) of the variables where P ⊂ V, and the edge weights 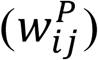 within this subset are greater than a predefined threshold. In this simulation, we selected 0.8, 0.6, 0.4, and 0.3 as the predefined thresholds for the purpose of illustrating the capability of the proposed approach to select the highly correlated variables. If SNP-SELECT includes at least one of the predefined highly correlated variables, the performance of the algorithm in selecting proxy variables among the highly correlated ones is considered fulfilled.

### Douglas-fir Breeding Populations

The Douglas-fir breeding population was established by the Ministry of Forests, Lands and Natural Resource of British Columbia, Canada in 1975 and consists of 165 full-sib families generated from structured paired-matings among 54 parents. The 1,372 individual trees used in this study consist of a subset of the full population and contains 37 full-sib families from 38 parents (see (54), for complete details). SNP genotypes for these 1,372 trees were generated using exome capture (55), resulting in 106,099 SNPs with missing ratio threshold less than 25% and minimum minor allele frequency (MAF) greater than 5% which comprises the ‘original’ data set.

The average numerator relationship matrix (***A***-matrix) of the DF dataset is known due to the structured pedigree, and was used as a baseline for comparison. We calculated the genomic estimated relatedness (***G-***matrix) using R package “rrBLUP” (56) using the mean imputation option on the original SNP dataset, as well as the five *k*-dominating SNP subsets. The pedigree-based relatedness (***A***-matrix) elements were compared to those of the ***G***-matrices of the selected subsets for validation.

### Grasshopper Mouse SNP Data

Grasshopper mouse (genus, *Onychomys*) are cricetid rodents that inhabit prairies, deserts and desert grasslands throughout the western United States, northern Mexico, and south-central Canada (57). Whereas *O. leucogaster* is readily distinguished based on body size, the two smaller species, *O. arenicola* and *O. torridus*, are morphologically cryptic and were treated as a single species until 1979 (58). The SNP dataset analyzed here was generated using genotyping-by-sequencing, GBS (19), as part of a study designed to test for evidence of hybridization at a site in southwestern New Mexico where all three species come into contact (59), and at other sites in New Mexico and Arizona where *O. leucogaster* is sympatric with *O. arenicola* and *O. torridus*, respectively. SNPs were called using a reference-free SNP discovery protocol (UNEAK pipeline (60)), and filtered based on a minor allele frequency (5%) and missing ratio (10%).

## Results

### Simulation Studies

When the SNP-SELECT algorithm was applied to the synthetic network with *k* ∈ {1,2,3,4,5}, all the *k*-dominating sets found included the predefined independent variables. The performance of the *k*-dominating set model in the selection of proxy variables is presented in Figure 2, with the predefined highly correlated variable thresholds *λ* ∈ {0.8,0.7,0.6,0.5,0.4,0.3}. As shown in Figure 2(a), the definition of highly correlated variables was strict 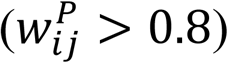 under this condition of few, highly connected variables, the use of larger values of either *k* or *λ* was encouraged. Also shown in Figure 2(b), 2(c) and 2(d), when relationships between variables are a mixture of high and low correlations, our results suggest the use of smaller values in *k* and *λ* to capture all relationships. By varying on *k* and *λ*, we demonstrate the flexibility and strength of SNP-SELECT in choosing proxy variables from highly correlated variables.

**Fig 2.**
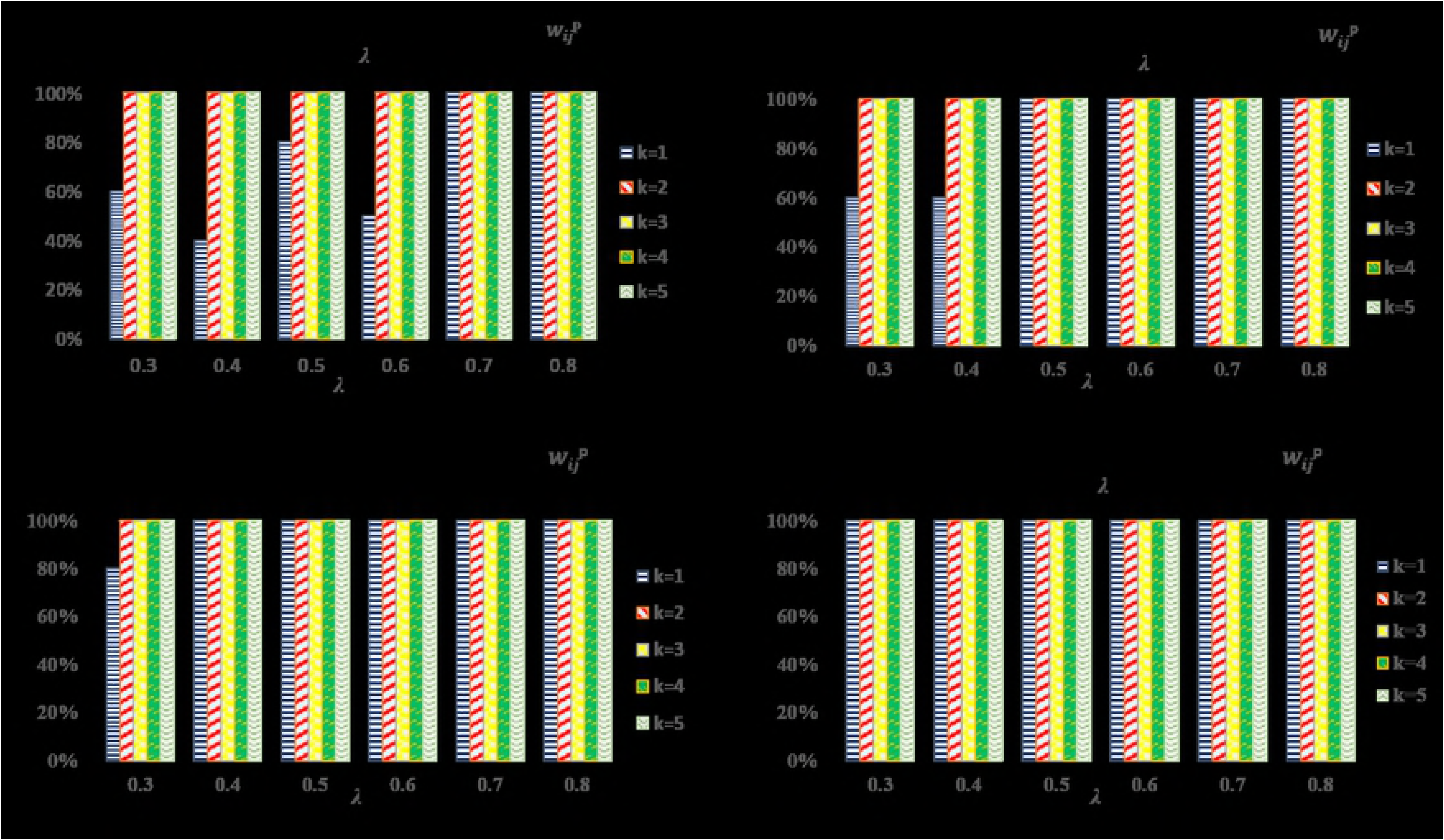
The capability of *k*-dominating set in selecting proxy variables among highly correlated variables. Ten synthetic undirected networks with *n* = 1,000 vertices (*V*) were simulated. (a) highly correlated variables defined as 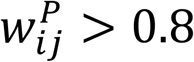; (b) highly correlated variables defined as 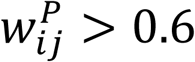 (c) highly correlated variables defined as 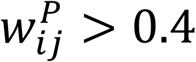 (d) highly correlated variables defined as 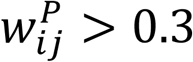.

### Pedigree Recovery for Douglas-Fir Breeding Populations

The SNP-SELECT algorithm was applied to select the influential SNPs to reconstruct the known pedigree for a Douglas-fir (DF) breeding population. Four *k*-dominating sets (DF107, DF105, DF103, DF102) with *k* = 1, and *λ* ∈ {0.7, 0.5, 0.3, 0.2} were generated. Among the four 1-dominating sets, DF103 has the best performance as shown in Table 1. To further investigate the impact of *k* on variable selection, another *k*-dominating set, DF203, with *k* = 2 and *λ* = 0.3 was generated to compare with DF103. The number of selected SNPs in DF107, DF105, DF103, DF102 and DF203 is 80,735, 67,062, 51,415, 41,539, and 68,188, respectively.

**Table 1:** The average difference of the upper triangle and the diagonal between pedigree-based relatedness (***A-***matrix) and genomic estimated relatedness (***G-***matrix). The best selected-subset for pedigree reconstruction (subset DF103) is highlighted. *λ* is linkage disequilibrium estimate.

| | $k$ | $\lambda$ | Num. of SNP | Ave. difference upper triangle | Ave. difference diagonal |
| --- | --- | --- | --- | --- | --- |
| Original Data | - | - | 106,099 | 0.034353 | 0.180374 |
| DF107 | 1 | 0.7 | 80,735 | 0.034240 | 0.103673 |
| DF105 | 1 | 0.5 | 67,062 | 0.034139 | 0.055994 |
| **DF103** | **1** | **0.3** | **51,415** | **0.034018** | **0.019769** |
| DF102 | 1 | 0.2 | 41,539 | 0.034249 | 0.123494 |
| DF203 | 2 | 0.3 | 68,188 | 0.034180 | 0.123950 |
| Random subset | - | - | 51,415 | 0.034498 | 0.180419 |
| COR03 | - | 0.3 | 39,768 | 0.034774 | 0.234326 |
| LRTag03 | - | 0.3 | 51,022 | 0.034292 | 0.135324 |

The performance of the five *k*-dominating sets generated better results than the original dataset for both average absolute upper triangle and diagonal differences from the genomic relationship matrix (***G***-matrix) to the traditional pedigree-based average numerator relationship matrix (***A***-matrix) (Table 1). Comparing the five *k*-dominating sets indicated that the DF103 subset performed best on pedigree recovery, especially for the diagonal pedigree information recovery. Figure 3 further illustrates the efficiency of the DF103 subset on pedigree reconstruction, and indicates that the ***G***-matrix generated from the DF103 subset was closer to the known ***A***-matrix as compared with the original dataset’s ***G***-matrix. Additionally, we randomly selected 10 subsets with the same SNP number as DF103 from the original dataset and used the average results of these 10 subsets to represent the performance of the randomly selected subset. The results indicated that all five *k*-dominating sets outperformed the randomly selected subset (Table 1).

**Fig 3:**
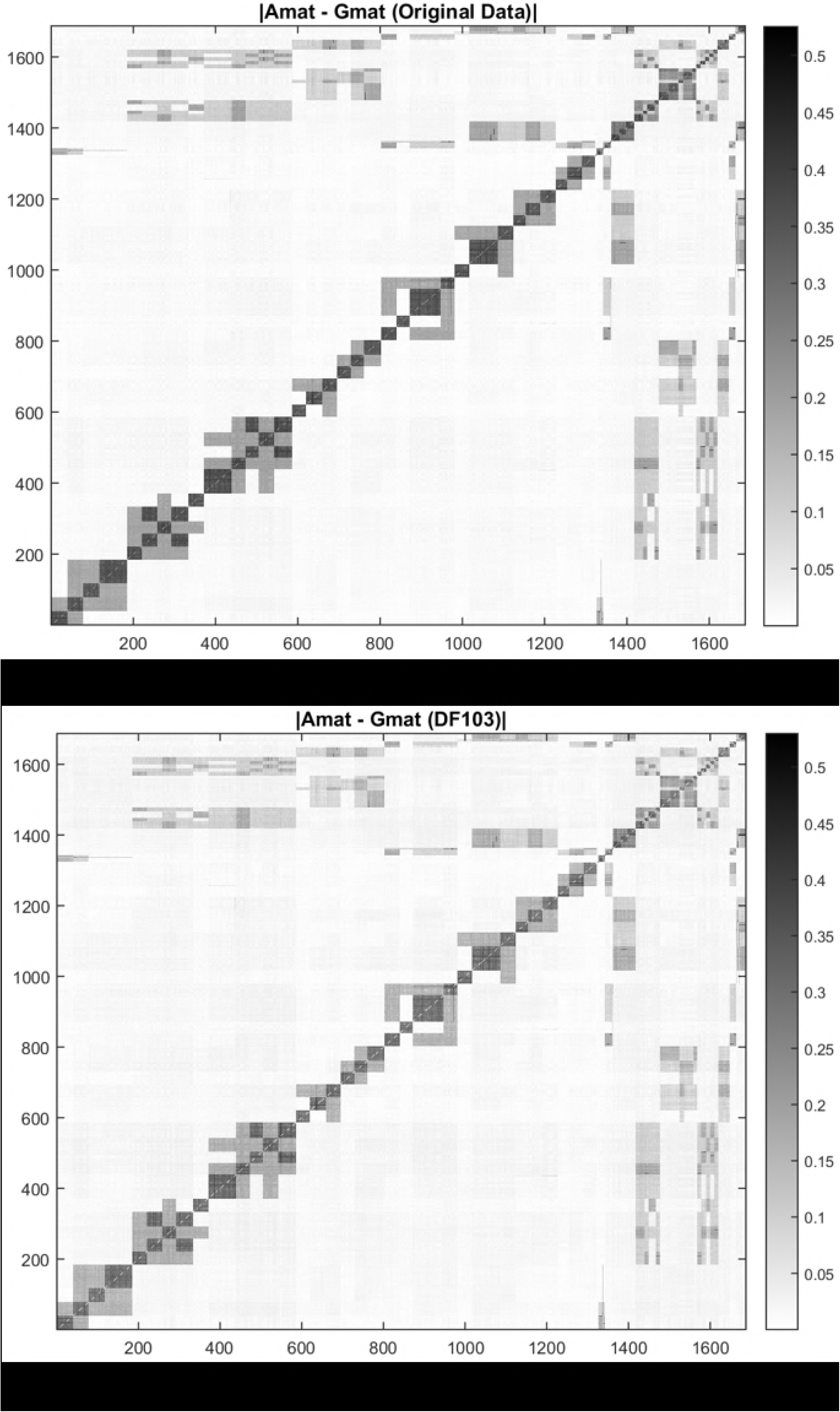
(a) Heatmap of the absolute difference between pedigree-based relatedness (***A***-matrix) and genomic estimated relatedness (***G***-matrix) generated from original data; (b) Heatmap of the absolute difference between pedigree-based relatedness (***A-***matrix) and genomic estimated relatedness (***G-***matrix) generated from DF103 subset. The color of Figure 3(b) is lighter than Figure 3(a). The lighter the color, the closer the relationship between ***A-***and ***G-***matrices of Douglas-fir breeding population.

The effectiveness of SNP-SELECT was also examined by the conventional approach that filters for SNP variables by pairwise correlation coefficients, as well as the LRTag algorithm that applies minimum set covering for SNP selection (61). The discrepancy between ***A-***and ***G-***matrices resulted from using correlation coefficient of 0.3 and *λ* = 0.3 was listed in Table 1 as COR03 and LRTag03, for pairwise correlation coefficient method and LRTag algorithm, respectively. Among all tests, the DF103 from SNP-SELECT remained the best SNP subset for estimating genetic relationship of Douglas-fir breeding population. Consider computing time requirement, when values of distance or pair-wise linkage disequilibria were pre-computed, SNP variable selection for SNP-SELECT could be complete in 8-10 minutes, while LRTag required about 18 hours for the same datasets.

### Clustering Analysis for Grasshopper Mouse Populations

To investigate parameters influencing population genetics of grasshopper mouse populations, 85,812 SNPs were used to genotype 226 samples representing three species: *O. arenicola*, (*n* = 76) *O. leucogaster*, (*n* = 67) and *O. torridus* (*n* = 83), collected at 12 geographic locations (Table 2). The dataset was pre-filtered based on a maximum of 10% missing data, and minimum MAF (minor allele frequency) of 5%. The SNP-SELECT was applied to generate three SNP subsets (MICE103, MICE105 and MICE107) with *k* = 1 and *λ* ∈ {0.3, 0.5, 0.7}, respectively; the number of informative SNPs retained in MICE103, MICE105 and MICE107 was 2,144, 11,014, and 22,355, respectively. The missing data in the original dataset and the three *k*-dominating sets was imputed with the most frequent genotype. Before the geographic origin analysis, we split the 226 samples into 3 groups based on species identity. There were 5 sampling locations in each species group (Table 2).

**Table 2:** Geographic location of grasshopper mouse (*Onychomys*) samples

| Species | Site Name (Sample Size) |
| --- | --- |
| *O. arenicola* | Animas/Rodeo, NM (20);<br>Pancho Villa, Chihuahua, Mex (7);<br>Organ Mountains, NM (27);<br>Sevilleta National Wildlife Refuge, NM (14);<br>Hidalgo del Parral, Chihuahua, Mex (8). |
| *O. leucogaster* | Petrified Forest, AZ (13);<br>Animas/Rodeo, NM (11);<br>Sevilleta National Wildlife Refuge, NM (14);<br>Felt, OK (19);<br>Garden City, KS (10). |
| *O. torridus* | Lone Pine, CA (11);<br>Carefree, AZ (8);<br>Santa Rita Experimental Range, AZ (19);<br>Animas/Rodeo, NM (28);<br>Chiricahua Mountains, AZ (17). |

The performance of the three *k*-dominating sets’ ability to predict the geographic origin of samples within each species was first evaluated using the *k-*means clustering approach in R (62). Clustering was initiated with *k* = 5, random seed at 20 and nstart = 100, where nstart specifies the initial configurations, and the algorithm will report on the best one (63, 64). The Adjusted Rand Index (ARI), a measure of agreement between clustering results and external criteria (65, 66), was used to evaluate the clustering results. As shown in Table 3, the clustering results for the largest SNP subset, MICE107, had the same performance as the original data of 85,812 SNPs in recovering the geographic origin of *O. arenicola* and *O. torridus* samples; however, MICE107 subset outperformed the original SNP data in recovering the geographic origin of *O. leucogaster* samples. Moreover, the clusters resulting from MICE105 and MICE103 exhibited larger ARI values than those from the original SNP data, indicative of a greater agreement and reduced errors in the clustering reached by SNP-SELECT variable selection. Overall, the MICE105 SNP subset (11,014 SNPs) demonstrated the greatest agreement among all selected subsets (Table 3).

**Table 3:** The adjusted rand index (ARI) shows the agreement between the computed clusters using *k*-means clustering algorithm and partitioning around medoids (PAM) algorithm with *k* = 5, using the original grasshopper mouse SNP data set and the *k*-dominating subsets. ARI values listed below show the agreement measurement between original sample locations and clustering results.

| Method | Dataset | SNP Number | *O. arenicola* | *O. leucogaster* | *O. torridus* |
| --- | --- | --- | --- | --- | --- |
| $k$-means | Original data | 85,812 | 0.3868 | 0.5981 | 0.5692 |
|  | MICE107 | 22,355 | 0.3868 | 0.7158 | 0.5692 |
|  | **MICE105** | **11,014** | **0.5256** | **0.7158** | **0.6003** |
|  | MICE103 | 2,144 | 0.3963 | 0.7158 | 0.6003 |
| PAM | Original data | 85,812 | 0.0706 | 0.2229 | 0.2244 |
|  | **MICE107** | 22,355 | 0.0513 | **0.3852** | 0.1812 |
|  | MICE105 | 11,014 | 0.1016 | 0.3509 | 0.2445 |
|  | **MICE103** | 2,144 | **0.1172** | 0.2902 | **0.2793** |

To confirm that the performance of SNP-SELECT was not the result of a specific clustering algorithm, the partitioning around medoids (PAM) algorithm (67) with *k* = 5 (random seed = 20) was performed using samples’ dissimilarity matrix of each species. To describe the dissimilarity matrix, we first define the ***G***-matrix as 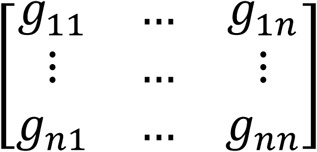. Then the dissimilarity matrix is defined as 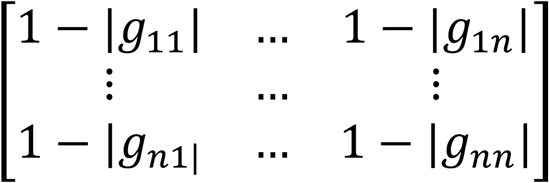. In Table 3, clusters resulting from the PAM algorithm also demonstrated that the selected subsets perform better than the original data in predicting actual sampling localities.

## Discussion

Owing to technological advancement in DNA sequencing methods, life scientists are grappling with exceedingly large data sets (68). The most obvious challenge is the large amount of genomic variation that needs to be processed and quantified in a very short time period (69). Although various data techniques have been adopted, the resulting data sets have several characteristics that make downstream analyses challenging (70). The common ones are: the number of variables is often much larger than the number of individuals, and data sets are usually sparse regarding relevant information, *i.e.* only a small subset of variables is associated with the phenotypic variation (71).

In genetic analyses using high dimensional data sets where there are more parameters than observations, penalized regression techniques are often required to ensure stable estimates (72, 73). The estimates of SNP marker effects are strongly affected by collinearity between predictors through a “grouping effect”-groups of variables highly correlated with other groups (of variables) sporadically (74). Such multicollinearity would further confound gene expression values obtained from DNA microarrays or determination of significance of SNP causality in genome-wide association (GWAS) or genomic selection (GS) studies (75-77). As a result, multiple-step GWAS analysis that includes SNP variable selection has been explored (16, 78, 79). While adoption of these methods might be an advantage when seeking functional variants associated with traits of interest, these fitness-associated SNP variables would bias inferences of gene flow, migration or dispersal (27, 80), and estimates of relatedness and inbreeding depression (81).

Without the dependency on phenotypes, SNP variable selection methods currently focus on pairwise correlations between variables (e.g. (82)). In principle, SNP variables are selected if the absolute value of a pairwise correlation (|*corr i, j* |) is less than a predefined threshold *λ*; or if |*corr i, j* | is no less than the given threshold, only the second variable will be selected (e.g. if |*corr i, j* | ≥ *λ*, SNP *j* will be selected). Here, we demonstrate the superior performance of the proposed *k*-dominating set variable selection over the conventional method of pairwise correlation coefficients (Table 1, COR03). As shown in Figure 3, diagonal values, indicative of the errors in estimating individuals’ genomic relationship based on markers, were minimized using SNP-SELECT. The pairwise estimates of genomic relationships (off-diagonal elements) were, however, mostly preserved (Table 1), suggesting that both the hidden and historical relatedness among individuals could still be recovered by the set of SNP variables selected by SNP-SELECT.

The use of genomic markers to uncover hidden relationships and potential pedigree errors in open-pollinated progeny has been effective in tree breeding programs (83, 84). Such pedigree reconstruction is a preferred method to determine the genealogical relationship among groups of related individuals, leading to improved estimation of genetic parameters (85-87). To maximize the advantage of using dense genomic markers, VanRaden (88) derived estimates of marker-based relationships between pairs of individuals as a genomic relationship matrix (***G***-matrix), which can be used as a substitute for the traditional pedigree-based average numerator relationship matrix (***A****-*matrix) in Henderson’s animal model (4, 89, 90). Also, combining the ***A***-matrix and the ***G***-matrix into a single genetic relationship matrix (***H***-matrix) has proven to be an effective approach to improve the relationship coefficients for better genetic parameter estimation (91, 92) and marker effect estimation (93), and to leverage extra phenotypic information from the non-genotyped individuals (90). To ensure improved accuracy in such single-step methods, the ***G****-* and ***A***-matrices should be compatible (94), and diagonal elements in the ***G****-*matrix need to be consistent with the ***A***-matrix diagonal elements; therefore rescaling ***A***- and ***G***-matrices would reflect the mean difference between these matrices (95), a context in which using SNP markers selected by SNP-SELECT could be considerably beneficial.

A C++ source code that implements SNP-SELECT and uses Gurobi(tm) optimization solver for the *k*-dominating set variable selection

## Conclusions

The *k*-dominating set model provides a flexible and effective method for selecting informative SNPs; a C++ source code (SNP-SELECT) that uses Gurobi^™^ Optimization solver is also released with the manuscript. This approach is scalable through the use of integer programming solvers and graph preprocessing, and can be extended to other genomic applications.

Using pedigree reconstruction and cluster analysis, the capacity of SNP-SELECT was demonstrated for solving the variable selection conundrum of large datasets without any significant runtime considerations. Furthermore, SNP-SELECT does not depend on the use of LD to define threshold for edges; other similarity/distance measure would broaden its applicability beyond breeding science and ecological genetics. Future work on the algorithmic aspects of this approach could focus on the development of graph and model decomposition techniques, as well as preprocessing techniques to improve scalability in practice.

## Declarations

### Availability of data and material

SNP-SELECT and SNP datasets used in this manuscript are available on the GitHub repository (https://github.com/transgenomicsosu/SNP-SELECT).

### Authors’ contributions

SS, ZM, BB and CC participated the design of the study and developed the method; SS and ZM implemented the algorithm and SS generated the simulated data, performed analysis, interpreted results; SS and CC drafted the manuscript. BR and YAE provided the Douglas-fir SNP data and SS, BR, YAE and CC analyzed Douglas-fir SNP dataset. SS, PC and BP examined SNP-SELECT’s performance on grasshopper mouse clustering analyses. All authors read and approved the final manuscript.

## Acknowledgements

We thank Michael Stoebr from the Ministry of Forests, Lands and Natural Resource Operations, British Columbia, for the use of Douglas-fir data. We are also grateful to Ashlee Rowe, and to the following institutions for providing grasshopper mouse tissue samples: Museum of Southwestern Biology, Museum of Vertebrate Zoology, Oklahoma State University Collection of Vertebrates. The project was completed with support from the High Performance Computing Center Facilities of Oklahoma State University.

## References

1. Song J, Carver BF, Powers C, Yan L, Klápště J, El-Kassaby YA, et al. Practical application of genomic selection in a doubled-haploid winter wheat breeding program. Mol Breed. 2017;37(10):117.

2. Poland J, Endelman J, Dawson J, Rutkoski J, Wu S, Manes Y, et al. Genomic selection in wheat breeding using genotyping-by-sequencing. The Plant Genome. 2012;5(3):103–13.

3. Heffner EL, Jannink J-L, Iwata H, Souza E, Sorrells ME. Genomic selection accuracy for grain quality traits in biparental wheat populations. Crop Sci. 2011;51(6):2597–606.

4. Habier D, Fernando RL, Garrick DJ. Genomic BLUP decoded: a Look into the black box of genomic prediction. Genetics. 2013;194(3):597–607.

5. Desta ZA, Ortiz R. Genomic selection: genome-wide prediction in plant improvement. Trends Plant Sci. 2014;19(9):592–601.

6. Bermingham M, Pong-Wong R, Spiliopoulou A, Hayward C, Rudan I, Campbell H, et al. Application of high-dimensional feature selection: evaluation for genomic prediction in man 2015. 10312 p.

7. Pes B, Dessì N, Angioni M. Exploiting the ensemble paradigm for stable feature selection: a case study on high-dimensional genomic data. Information Fusion. 2017;35:132–47.

8. He Q, Lin D-Y. A variable selection method for genome-wide association studies. Biometrics. 2011;27(1):1–8.

9. Ayers KL, Cordell HJ. SNP selection in genome-wide and candidate gene studies via penalized logistic regression. Genet Epidemiol. 2010;34(8):879–91.

10. Tibshirani R. Regression shrinkage and selection via the Lasso. J R Stat Soc Series B Stat Methodol. 1996;58:267–88.

11. Fan J, Lv J. A selective overview of variable selection in high dimensional feature space Statistica Sinica. 2010;20(1):101.

12. Mehmood T, Liland KH, Snipen L, Sæbø S. A review of variable selection methods in Partial Least Squares Regression. Chemometrics Intellig Lab Syst. 2012;118:62–9.

13. Fan J, Lv J. Sure independence screening for ultrahigh dimensional feature space. J Roy Stat Soc Ser B (Stat Method). 2008;70(5):849–911.

14. Wasserman L, Roeder K. High dimensional variable selection. Annals of statistics. 2009;37(5A):2178–201.

15. Bogdan M, van den Berg E, Sabatti C, Su W, Candès EJ. SLOPE—adaptive variable selection via convex optimization. The annals of applied statistics. 2015;9(3):1103–40.

16. Dehman A, Ambroise C, Neuvial P. Performance of a blockwise approach in variable selection using linkage disequilibrium information. BMC Bioinformatics. 2015;16(1):148.

17. Luikart G, England PR, Tallmon D, Jordan S, Taberlet P. The power and promise of population genomics: from genotyping to genome typing. Nat Rev Genet. 2003;4:981.

18. Hohenlohe PA, Bassham S, Etter PD, Stiffler N, Johnson EA, Cresko WA. Population genomics of parallel adaptation in threespine stickleback using sequenced RAD tags. PloS Genet. 2010;6(2):e1000862.

19. Elshire RJ, Glaubitz JC, Sun Q, Poland JA, Kawamoto K, Buckler ES, et al. A Robust, Simple Genotyping-by-Sequencing (GBS) Approach for High Diversity Species. PloS ONE. 2011;6(5):e19379.

20. Chen C, Mitchell SE, Elshire RJ, Buckler ES, El-Kassaby YA. Mining conifers’ mega-genome using rapid and efficient multiplexed high-throughput genotyping-by-sequencing (GBS) SNP discovery platform. Tree Genetics & Genomes. 2013;9(6):1537–44.

21. Bonhomme M, Chevalet C, Servin B, Boitard S, Abdallah J, Blott S, et al. Detecting selection in population trees: The lewontin and krakauer test extended. Genetics. 2010;186(1):241–62.

22. Beaumont MA, Balding DJ. Identifying adaptive genetic divergence among populations from genome scans. Mol Ecol. 2004;13(4):969–80.

23. Foll M, Gaggiotti O. A genome-scan method to identify selected loci appropriate for both dominant and codominant markers: A bayesian perspective. Genetics. 2008;180(2):977–93.

24. Guo F, Dey DK, Holsinger KE. A bayesian hierarchical model for analysis of Single-Nucleotide Polymorphisms diversity in multilocus, multipopulation samples. Journal of the American Statistical Association. 2009;104(485):142–54.

25. Vitti JJ, Grossman SR, Sabeti PC. Detecting natural selection in genomic data. Annu Rev Genet. 2013;47(1):97–120.

26. Nielsen R. Statistical tests of selective neutrality in the age of genomics. Heredity. 2001;86:641.

27. Kirk H, Freeland JR. Applications and implications of neutral versus non-neutral markers in Molecular Ecology. Int J Mol Sci. 2011;12(6):3966.

28. Excoffier L, Dupanloup I, Huerta-Sánchez E, Sousa VC, Foll M. Robust demographic inference from genomic and SNP data. PloS Genet. 2013;9(10):e1003905.

29. Robertson A. Gene frequency distributions as a test of selective neutrality. Genetics. 1975;81(4):775–85.

30. Jain AK, Dubes RC. Algorithms for clustering data: Prentice-Hall, Inc.; 1988.

31. Jambu M, Lebeaux M-O. Cluster analysis and data analysis: Elsevier Science Ltd; 1983.

32. Spath H. Cluster analysis algorithms for data reduction and classification of objects: Chichester: Ellis Horwood; 1980.

33. West DB. Introduction to graph theory: Prentice hall Upper Saddle River; 2001.

34. Diestel R. Graph Theory. 5 ed: Springer-Verlag Berlin Heidelberg; 2017. XVIII, 429 p.

35. Haynes TW, Hedetniemi S, Slater P. Fundamentals of domination in graphs: Marcel Dekker Inc.; 1998.

36. White JG, Southgate E, Thomson JN, Brenner S. The structure of the nervous system of the nematode caenorhabditis elegans. Philosophical Transactions of the Royal Society of London Series B. 1986;314:1–340.

37. Watts DJ, Strogatz SH. Collective dynamics of ‘small-world’ networks. Nature. 1998;393:440.

38. Balasundaram B, Butenko S. Graph domination, coloring and cliques in telecommunications. In: Resende MGC, Pardalos PM, editors. Handbook of Optimization in Telecommunications. Boston, MA: Springer US; 2006. p. 865–90.

39. Michael RG, David SJ. Computers and intractability: a guide to the theory of NP-completeness: WH Free. Co., San Fr; 1979. 90–1 p.

40. Butenko S, Cheng X, Oliveira CA, Pardalos PM. A new heuristic for the minimum connected dominating set problem on ad hoc wireless networks. In: Butenko S, Murphey R, Pardalos PM, editors. Recent Developments in Cooperative Control and Optimization. Boston, MA: Springer US; 2004. p. 61–73.

41. Wolsey LA. Integer Programming: Wiley; 1998.

42. Wang C, Kao W-H, Hsiao CK. Using hamming distance as information for SNP-sets clustering and testing in disease association studies. PloS One. 2015;10(8):e0135918.

43. Bartlett CW, Yeon Cheong S, Hou L, Paquette J, Yee Lum P, Jäger G, et al. An eQTL biological data visualization challenge and approaches from the visualization community. BMC Bioinformatics. 2012;13(8):S8.

44. Zhang X, Pan F, Xie Y, Zou F, Wang W, editors. COE: a general approach for efficient genome-wide two-locus epistasis test in disease association study 2009; Berlin, Heidelberg: Springer Berlin Heidelberg.

45. vonHoldt BM, Pollinger JP, Lohmueller KE, Han E, Parker HG, Quignon P, et al. Genome-wide SNP and haplotype analyses reveal a rich history underlying dog domestication. Nature. 2010;464:898.

46. Shriver MD, Kennedy GC, Parra EJ, Lawson HA, Sonpar V, Huang J, et al. The genomic distribution of population substructure in four populations using 8,525 autosomal SNPs. Human Genomics. 2004;1(4):274.

47. Yang J, Benyamin B, McEvoy BP, Gordon S, Henders AK, Nyholt DR, et al. Common SNPs explain a large proportion of the heritability for human height. Nat Genet. 2010;42:565.

48. Liu G, Wang Y, Wong L. FastTagger: an efficient algorithm for genome-wide tag SNP selection using multi-marker linkage disequilibrium. BMC Bioinformatics. 2010;11(1):66.

49. Weng L, Macciardi F, Subramanian A, Guffanti G, Potkin SG, Yu Z, et al. SNP-based pathway enrichment analysis for genome-wide association studies. BMC Bioinformatics. 2011;12(1):99.

50. González-Martínez SC, Grivet D. Association mapping in plants. Oraguzie NC, Rikkerink EHA, Gardiner SE, Silva HND, editors: Springer, New York, NY; 2009. ix–x p.

51. Excoffier L, Slatkin M. Maximum-likelihood estimation of molecular haplotype frequencies in a diploid population. Mol Biol Evol. 1995;12(5):921–7.

52. Gurobi Optimization I. Gurobi optimizer reference manual 2018 [Available from: http://www.gurobi.com.

53. Chen C, DeClerck G, Tian F, Spooner W, McCouch S, Buckler E. PICARA, an analytical pipeline providing probabilistic inference about a priori candidates genes underlying genome-wide association QTL in plants. PloS One. 2012;7(11):e46596.

54. Thistlethwaite FR, Ratcliffe B, Klápště J, Porth I, Chen C, Stoehr MU, et al. Genomic prediction accuracies in space and time for height and wood density of Douglas-fir using exome capture as the genotyping platform. BMC Genomics. 2017;18(1):930.

55. Neves LG, Davis JM, Barbazuk WB, Kirst M. Whole-exome targeted sequencing of the uncharacterized pine genome. The Plant Journal. 2013;75(1):146–56.

56. Endelman JB. Ridge Regression and Other Kernels for Genomic Selection with R Package rrBLUP. The Plant Genome. 2011;4(3):250–5.

57. Hall ER. The mammals of North America. second ed: John Wiley and Sons, New York; 1981.

58. Hinesley LL. Systematics and distribution of two chromosome forms in the southern grasshopper mouse, genus onychomys. J Mammal. 1979;60(1):117–28.

59. Sullivan RM, Hafner DJ, Yates TL. Genetics of a contact zone between three chromosomal forms of the grasshopper mouse (genus onychomys): A reassessment. J Mammal. 1986;67(4):640–59.

60. Lu F, Lipka AE, Glaubitz J, Elshire R, Cherney JH, Casler MD, et al. Switchgrass Genomic Diversity, Ploidy, and Evolution: Novel Insights from a Network-Based SNP Discovery Protocol. PloS Genetics. 2013;9(1):e1003215.

61. Liu L, Wu Y, Lonardi S, Jiang T. Efficient genome-wide TagSNP selection across populations via the linkage disequilibrium criterion. Journal of Computational Biology. 2010;17(1):21–37.

62. Team RDC. R: A Language and Environment for Statistical Computing. 2011.

63. Muca M, Kutrolli G, Kutrolli M. A proposed algorithm for determining the optimal number of clusters. European Scientific Journal, ESJ. 2015;11(36).

64. Jay JJ, Eblen JD, Zhang Y, Benson M, Perkins AD, Saxton AM, et al. A systematic comparison of genome-scale clustering algorithms. BMC Bioinformatics. 2012;13(10):S7.

65. Yeung KY, Ruzzo W. Details of the adjusted rand index and clustering algorithms supplement to the paper “An empirical study on Principal Component Analysis for clustering gene expression data” (to appear in Bioinformatics)2001.

66. Santos JM, Embrechts M, editors. On the use of the adjusted rand index as a metric for evaluating supervised classification 2009; Berlin, Heidelberg: Springer Berlin Heidelberg.

67. Maechler M, Rousseeuw P, Struyf A, Hubert M, Hornik K. cluster: cluster analysis basics and extensions. 2017.

68. Marx V. The big challenges of big data. Nature. 2013;498:255.

69. May M. Life science techologies: big biological impacts from big data. Science. 2014;344(6189):1298–300.

70. Yang Y, Han L, Yuan Y, Li J, Hei N, Liang H. Gene co-expression network analysis reveals common system-level properties of prognostic genes across cancer types. 2014;5:3231.

71. Degenhardt F, Seifert S, Szymczak S. Evaluation of variable selection methods for random forests and omics data sets. Brief Bioinform. 2017:bbx124–bbx.

72. Meuwissen THE, Hayes BJ, Goddard ME. Prediction of total genetic value using genome-wide dense marker maps. Genetics. 2001;157(4):1819–29.

73. Hastie T, Tibshirani R, Friedman J. The elements of statistical learning: Springer series in statistics New York; 2001.

74. Wimmer V, Lehermeier C, Albrecht T, Auinger H-J, Wang Y, Schön C-C. Genome-wide prediction of traits with different genetic architecture through efficient variable selection. Genetics. 2013;195(2):573–87.

75. Hong S, Kim Y, Park T. Practical issues in screening and variable selection in genome-wide association analysis. Cancer Inform. 2014;13(Suppl 7):55–65.

76. Ishwaran H, Rao JS. Geometry and properties of generalized ridge regression in high dimensions. Contemp Math. 2014;622:81–93.

77. El-Kassaby YA. Associations between allozyme genotypes and quantitative traits in Douglas-fir [Pseudotsuga menziesii (Mirb.) Franco]. Genetics. 1982;101(1):103–15.

78. Cho S, Kim K, Kim YJ, Lee J-K, Cho YS, Lee J-Y, et al. Joint identification of multiple genetic variants via elastic-net variable selection in a genome-wide association analysis. Ann Hum Genet. 2010;74(5):416–28.

79. Szymczak S, Holzinger E, Dasgupta A, Malley JD, Molloy AM, Mills JL, et al. r2VIM: a new variable selection method for random forests in genome-wide association studies. BioData Min. 2016;9(1):7.

80. Holderegger R, Kamm U, Gugerli F. Adaptive vs. neutral genetic diversity: implications for landscape genetics. Landscape Ecol. 2006;21(6):797–807.

81. Chelo IM, Carvalho S, Roque M, Proulx SR, Teotónio H. The genetic basis and experimental evolution of inbreeding depression in Caenorhabditis elegans. Heredity. 2013;112:248.

82. Hainke K, Szugat S, Fried R, Rahnenführer J. Variable selection for disease progression models: methods for oncogenetic trees and application to cancer and HIV. BMC Bioinformatics. 2017;18(1):358.

83. Wang J. Sibship reconstruction from genetic data with typing errors. Genetics. 2004;166(4):1963–79.

84. Kalinowski ST, Taper ML, Marshall TC. Revising how the computer program cervus accommodates genotyping error increases success in paternity assignment. Mol Ecol. 2007;16(5):1099–106.

85. El-Kassaby YA, LstibŮRek M. Breeding without breeding. Genetics Research. 2009;91(2):111–20.

86. Klápště J, Lstibůrek M, El-Kassaby YA. Estimates of genetic parameters and breeding values from western larch open-pollinated families using marker-based relationship. Tree Genet Genom. 2014;10(2):241–9.

87. El-Kassaby YA, Cappa EP, Liewlaksaneeyanawin C, Klapste J, Lstiburek M. Breeding without breeding: is a complete pedigree necessary for efficient breeding? PLoS One. 2011;6(10):e25737.

88. VanRaden PM. Efficient Methods to Compute Genomic Predictions. Journal of Dairy Science. 2008;91(11):4414–23.

89. Henderson C. Applications of linear models in animal breeding. University of Guelph Press, Guelph. 1984;11:652–3.

90. Ratcliffe B, Gamal El-Dien O, Cappa EP, Porth I, Klapste J, Chen C, et al. Single-step BLUP with varying genotyping effort in open-pollinated Picea glauca. G3: Genes|Genomes|Genetics. 2017.

91. Legarra A, Aguilar I, Misztal I. A relationship matrix including full pedigree and genomic information. J Dairy Sci. 92(9):4656–63.

92. Misztal I, Legarra A, Aguilar I. Computing procedures for genetic evaluation including phenotypic, full pedigree, and genomic information. J Dairy Sci. 92(9):4648–55.

93. Wang H, Misztal I, Aguilar I, Legarra A, Muir WM. Genome-wide association mapping including phenotypes from relatives without genotypes. Genetics Research. 2012;94(2):73–83.

94. Christensen OF, Madsen P, Nielsen B, Ostersen T, Su G. Single-step methods for genomic evaluation in pigs. Animal. 2012;6(10):1565–71.

95. Powell JE, Visscher PM, Goddard ME. Reconciling the analysis of IBD and IBS in complex trait studies. Nature Reviews Genetics. 2010;11:800.

